# Is the distribution of plasmid lengths bimodal?

**DOI:** 10.1101/2023.09.10.557055

**Authors:** Ian Dewan, Hildegard Uecker

## Abstract

The length of a plasmid is a key property which is linked to many aspects of plasmid biology. When distributions of plasmid lengths are shown in the literature, they are usually plotted with length on a logarithmic scale. However, a quantity and its logarithm have distinct distributions which may differ considerably in shape. Mistaking the distribution of log-lengths for the distribution of lengths can therefore lead to distorted conclusions about the distribution; in particular, the distribution of log-lengths may be bimodal when the distribution of lengths is only unimodal. This particular confusion has arisen in the literature where the length distribution is often claimed to be bimodal based on examination of what is in fact the log-length distribution. While the length distribution is indeed bimodal within many bacterial families, it is not across the ensemble of all plasmids. We suggest that authors should be careful to show the plasmid length distribution, or to distinguish the two distributions, to avoid misleading inferences.

**Highlights:**

- The distributions of lengths and log-lengths of plasmids are different, and have considerably different shapes.
- The typical practice of using a logarithmic scale for plasmid lengths leads to confusion between the two distributions.
- In particular, the distribution of log-lengths can be bimodal when the distribution of lengths is not.
- The length distribution within bacterial families is often bimodal, but across all plasmids, it is unimodal.
- Clearly distinguishing between the distributions will ensure that biological conclusions drawn from them are robust.

## 1 Introduction

Bacterial plasmids have a wide variety of sizes, from tiny cryptic plasmids less than a kilobase long, to multiple-megabase megaplasmids that rival the size of chromosomes (Ciok et al. 2016; Hall et al. 2021). The length is closely linked to other aspects of a plasmid’s biology, since smaller plasmids contain fewer genes and may therefore be limited in their functions (plasmid-specific or related to the host phenotype), while very large plasmids may contain a wide variety of accessory genes.

It has generally been assumed in the literature that plasmids can be broadly divided into two groups on the basis of length, and that this division corresponds to a biologically meaningful distinction between two plasmid lifestyles. The shorter group, with a modal length less than 10 kbp (Hall et al. 2021), is said to comprise plasmids with high copy numbers which are mostly mobilizable or nontransmissible, while the longer group, with a modal length around 100 kbp, comprises low copy number plasmids which are more likely to be conjugative. Part of the evidence for the existence of two length-based groups is the distribution of plasmid lengths estimated using standard techniques (kernel density estimation or histograms): the resulting density estimates clearly have a bimodal shape (this is done for all plasmids together by Smillie et al. 2010; Garcillán-Barcia et al. 2023 and for plasmids divided by host family by Hall et al. 2021; Rodrí guez-Beltrán et al. 2021).

These graphs show densities with a logarithmic scale on the *x*-axis, which is a natural choice given that the lengths of plasmids extend over four orders of magnitude: however, when we plotted plasmid length data (pooled across bacterial families) on a linear scale (Dewan and Uecker 2023), the bimodal distribution disappeared (see also diCenzo and Finan 2017 who also use a linear scale and find a unimodal distribution). In this letter, we explain the reason for this discrepancy, and what implications, if any, it has for our understanding of plasmids.

## 2 Analysis and Results

### The distribution of log-lengths is not the distribution of lengths

Why are there two types of figure that make very different statements about the plasmid length distribution in the literature? They are in fact showing two different distributions: the actual distribution of lengths (shown in Figure 1A) and what, despite often being called the distribution of lengths, is in fact the distribution of *log-lengths* (shown in Figure 1C). In other words, the graphs with a log-scaled *x*-axis do not simply plot the same numerical values from the distribution of plasmid length but on a log scale (which would instead produce Figure 1B), but a distinct distribution. The distributions of plasmid lengths and log-lengths may have quite different shapes; in general, the shape of a distribution changes, possibly drastically, when a transformation of the random variable is applied. Intuitively, the reason for this change is that bins of equal sizes on the log scale correspond to bins of different sizes on the linear scale: a bin at small plasmid log-lengths contains a much narrower range of plasmid lengths than a bin at large plasmid log-lengths, and the latter might contain more sequences than the former for this reason alone. As we show in the next paragraph, to obtain the density of log-lengths, we need to multiply the distribution of lengths by an exponentially increasing function. This can convert a heavy tail of the distribution on the linear scale into an additional mode on the log scale. This is illustrated in Figure 2 by a hypothetical example of a distribution of a random variable and the corresponding distribution of its logarithm. The distribution of the variable on the linear scale is unimodal, but the distribution of its logarithm is bimodal, precisely what happens with plasmid lengths.

**Figure 1:**
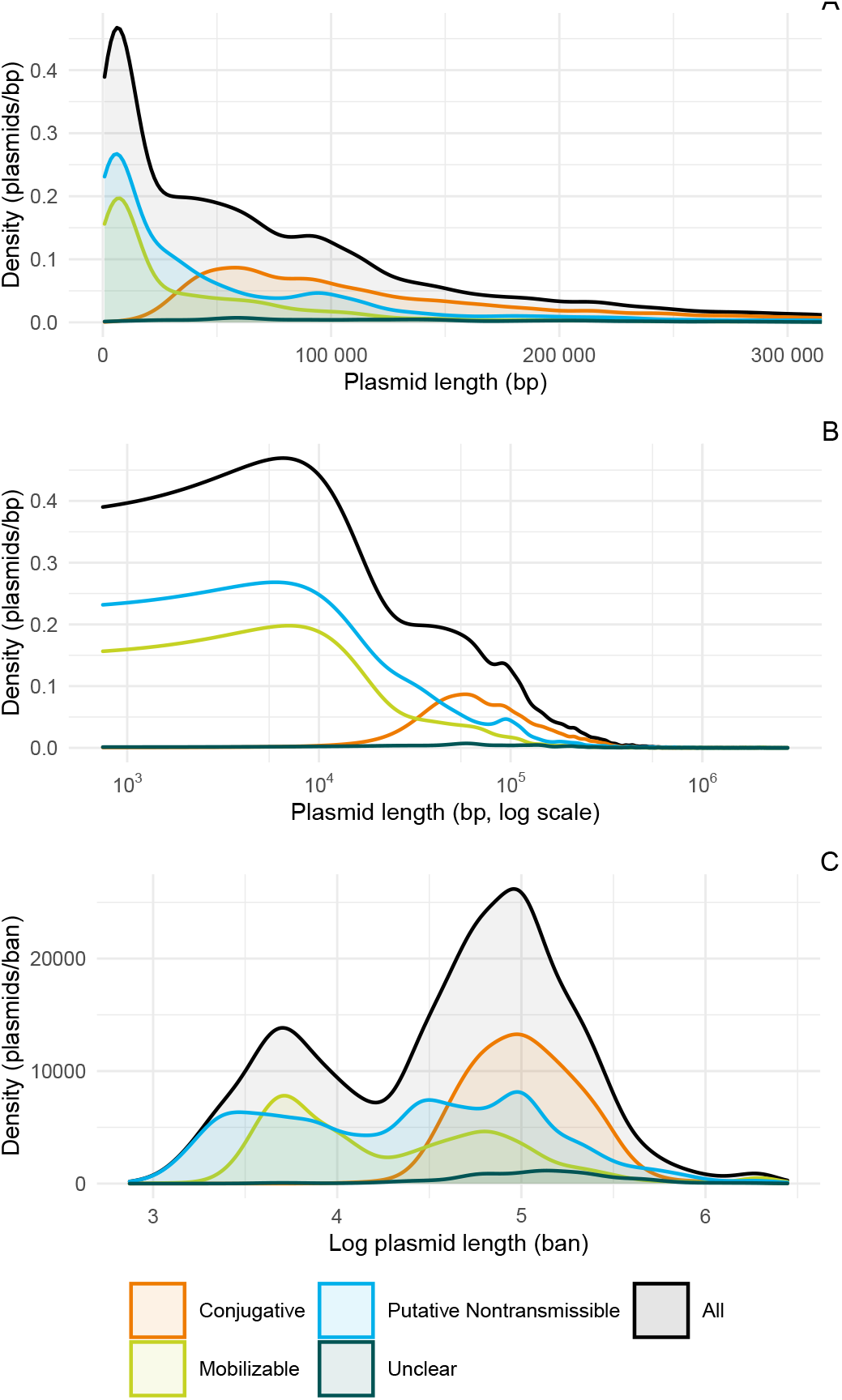
The distributions of plasmid lengths and log-lengths for all plasmids together and for each mobility class individually. The data is taken from PLSDB (Schmartz et al. 2021, retrieved 1 March 2023), and the distribution estimate made by kernel density estimation in R (R Core Team 2020). Panel A shows the estimated distribution of plasmid lengths (with length on a linear scale). Only plasmids of length less than 300 000 bp are shown for reasons of scale: 1844 plasmids in the database (5.34 % of the total) are longer than the limit. Based on Figure 1 of Dewan and Uecker (2023). Panel B shows the plasmid length distribution, estimated as in panel A, but with a log-scaled x-axis; i.e. kernel density estimation is performed on a linear scale, and the logarithm is only applied for the plot itself. Panel C shows the estimated distribution of plasmid log-lengths (“ban” are the units of the base-10 logarithm); i.e. the logarithm is applied before performing the density kernel estimation. To classify the plasmids by mobility, the annotations in PLSDB from MOB-typer (Robertson and Nash 2018), indicating putative relaxases or conjugative genes, have been used. Those plasmids with both are conjugative, those with only a relaxase are mobilizable, and those with neither are putatively nontransmissible (although Ares-Arroyo et al. 2022 have shown that many plasmids lacking a relaxase still have an *oriT*, and are therefore in fact mobilizable). The “unclear” class includes those plasmids which had conjugative genes but no relaxase. It is possible that these are misidentified conjugative plasmids, which have an unknown relaxase, or misidentified mobilizable or nontransmissible plasmids, which do not actually have conjugative genes.

**Figure 2:**
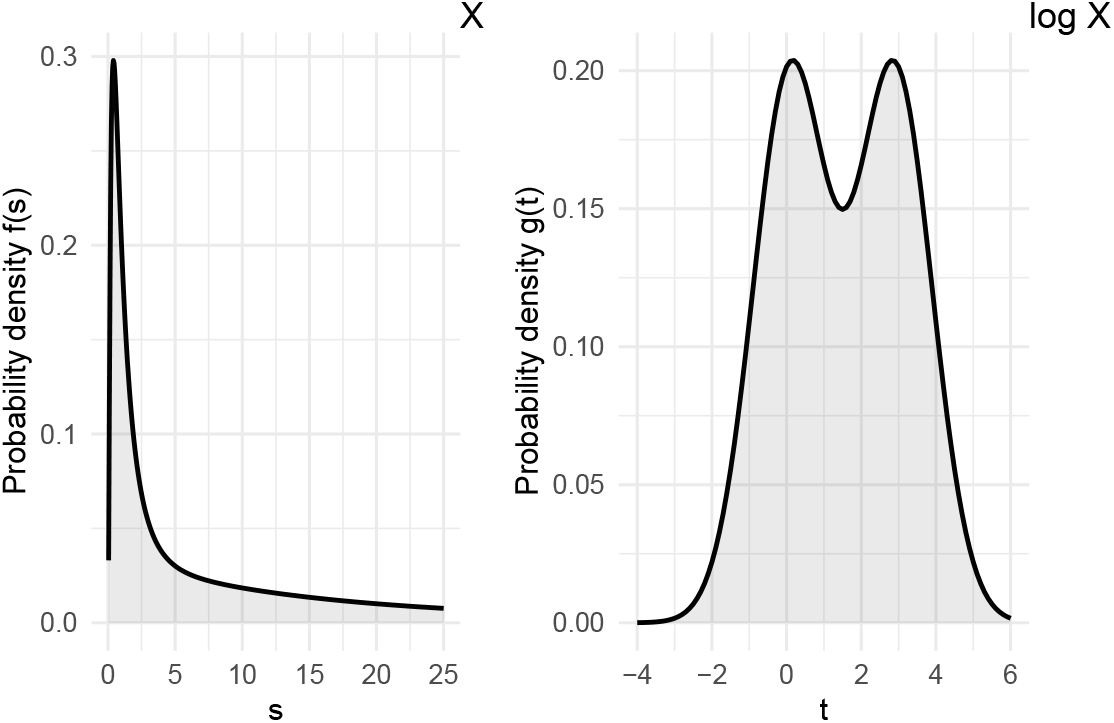
The probability density function of a distribution (left) and its (natural) logarithm (right). The distribution of log *X* is an equal mixture of two normal distributions, with means 0.1 and 2.9 and standard deviation 1; the distribution of *X* is therefore a mixture of two log-normal distributions.

We can show the change in the shape of the distribution formally by exam-ining the effect of a log transformation on the density function of a distribution. Suppose we have a (positive) random variable *X* with a probability density func-tion *f*. The defining characteristic of the probability density function is that it can be integrated over an interval to find the probability that *X* lies in that interval; that is for any numbers *a ≤ b*,

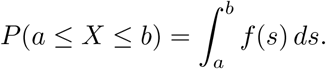

If we know *f*, we can find the probability density function of log *X*:

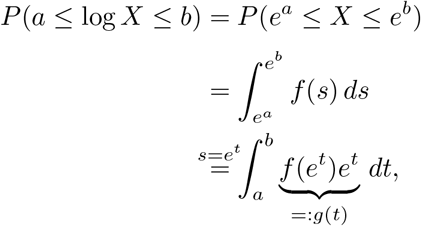

so the probability density function of log *X* is *g*(*t*) = *f* (*e*^*t*^)*e*^*t*^. (Here we have used the natural logarithm, but the same argument applies to the “common” or base-10 logarithm usually used in log scales.) When we apply the change of variable *s* = *e*^*t*^ in the above integral to pass from integrating on a linear scale to integrating on a log scale, we multiply *f* by a nonconstant—indeed exponentially increasing—function *e*^*t*^, changing the shape of the function. If the probability density function *f* decreases too slowly (i.e. if it has a heavy tail), this creates a second mode.

The difference in the number of modes between Figure 1A and 1C is therefore simply a mathematical consequence of the log scale. Changes in the shape of the distribution of course also occur in subsets of plasmids as well, such as the distributions within mobility classes shown in Figure 1. Note, for example, that looking at the log-length distributions in Figure 1C would suggest that mobi-lizable plasmids have a bimodal distribution and nontransmissible plasmids are roughly equally abundant over a wide range of lengths; but the length distributions in Figure 1A show that both groups have a unimodal distribution with a long right tail.

To verify that these changes in distribution shape are indeed caused by the log transformation and are not instead artifacts of the mechanism of density estimation, we have confirmed that the general shape of the length and log-length distributions is consistent under different estimation methods: different bandwidth selections for kernel density estimation (results shown in supplementary figures S1 and S4), the empirical cumulative distribution function (shown in figures S2 and S5), and equal-area histograms (shown in figure S3). These do indeed all give similar results.

### Within bacterial families, the plasmid length distribution is often bimodal, but the modes and the valley are incorrectly predicted by the log-length distribution

Above we considered the plasmid length distribution for all plasmids together, irrespective of their bacterial hosts. However, the size of plasmids differs between groups of bacteria, for example because bacteria with larger genome sizes in general have larger plasmids (Slater et al. 2008; Smillie et al. 2010). If the length distribution is really bimodal within groups, but the modes (and the antimode, or local minimum density, between them) have different locations in different groups because of a general difference in size, then in the pooled dis-tribution for all plasmids the modes and antimodes of different groups might obscure each other. Looking at the length distribution by family, as shown in Figure 3, shows that some bacterial families indeed show a pronounced bimodal distribution on the linear scale (plasmids from Enterobacteriaceae, Enterococ-caceae, or Staphylococcaceae), while for others there is no clear second mode (plasmids from Bacillaceae or Moraxcellaceae), or it is very weak (plasmids from Lactobacillaceae). A caveat to the within family density estimates is that they are based on much fewer sequences than the estimate for all plasmids. We would like to point out that we have identified modes by eye, only counting what we visually identify as major modes and not taking any smaller secondary modes into account (which may or may not be due to sampling effects or artifacts of the kernel density estimation). In particular, we do not perform any statistical analysis to determine which of the small modes are “real” (using, e.g. Silverman 1981), which is not necessary to make the point of the article.

**Figure 3:**
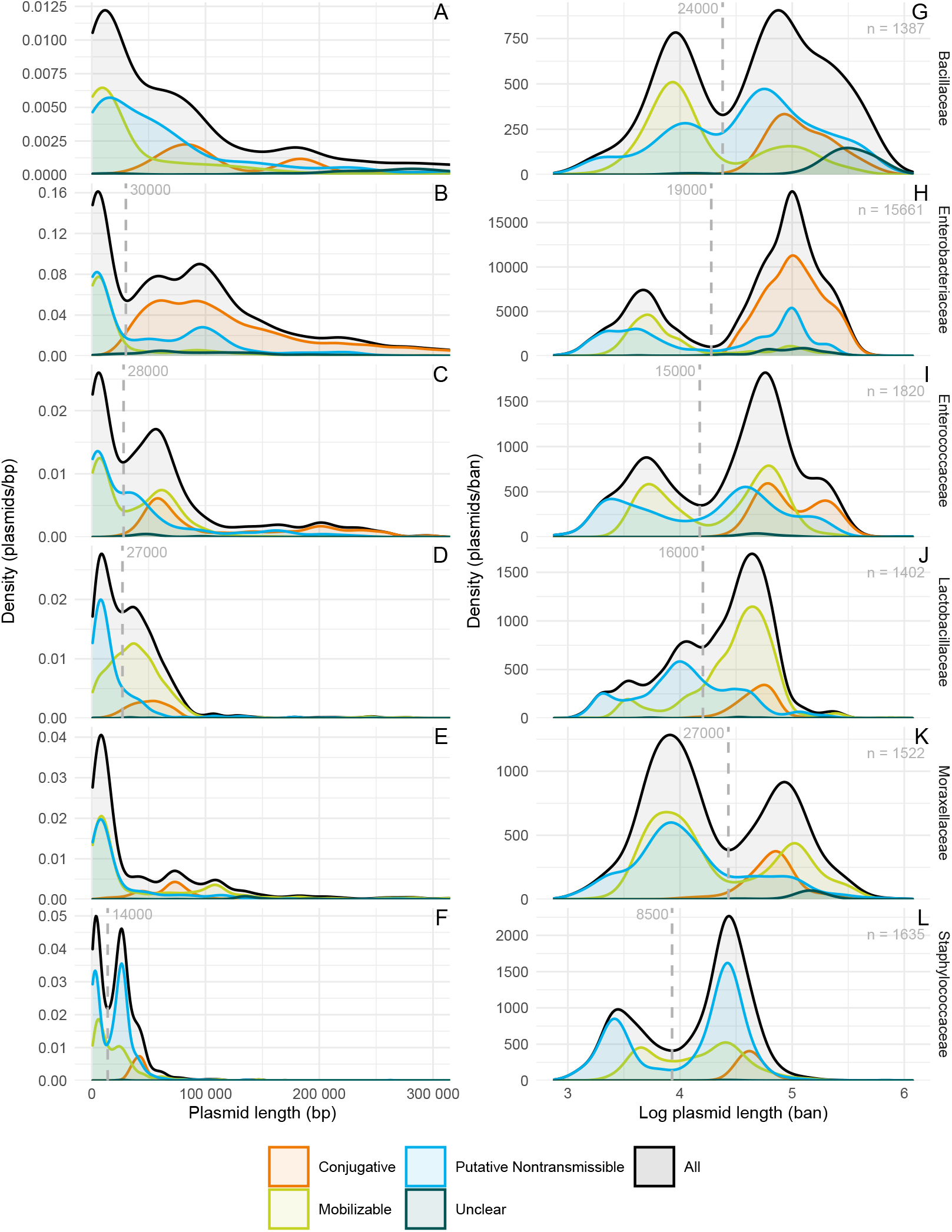
Distributions of plasmid length (left column, panels A–F) and loglength (right column, panels G–L) for plasmids from the six bacterial families with at least 1000 sequences in the database. Note that the length distributions are cut off at 300 000 bp for reasons of scale. The vertical lines show the location of the antimode of each distribution to the nearest kilobase. “Ban” are the units of the base-10 logarithm. The sample size for each family is given by *n* in the upper right corner of each row. For the data source and details on the figure construction see Figure 1.

Nevertheless, even if both distributions have the same number of modes, looking at the log-length distribution can be misleading, since taking logarithms can change not only the number but also the location of modes (and of antimodes). This shift in the mode can, for example, be seen for conjugative plasmids in Figure 1. The modal log-length of a conjugative plasmid appears to be about 5 (corresponding to a length of 10^5^ bp), while the modal length is smaller (closer to 50 000 bp). Similarly, within bacterial families where both the length and log-length distributions are bimodal, the antimode dividing small and large plasmids is not the same for the length and the log-length distributions (Fig-ure3). Thus even if there are distinct groups of “large” and “small” plasmids, trying to read their typical sizes or the dividing line between them off of an estimate of the distribution of log-lengths will be misleading. The accuracy of the location of modes and antimodes is, of course, in either case affected by the quality of the identification of plasmids in the database.

## 3 Discussion

We have shown that estimates of the distribution of plasmid lengths from available plasmid sequence data are not bimodal, at the global level at least, even though estimates of the distribution of log-lengths are, and that this kind of discrepancy is an ordinary feature of probability distributions. Within bacterial families, such a bimodal length distribution often exists, but the modes and antimode of the length distribution are incorrectly predicted by the log-length distribution. It is important to note that this does not mean that considering the distribution of log-lengths is wrong. There may well be reasons to be interested in the distribution of log-lengths rather than lengths. But the distribution of loglengths is different from the distribution of lengths, and information about the distribution of lengths cannot be directly read off the graph of the distribution of log-lengths.

How did the confusion of the length and log-length distributions arise? We suspect that it is due to combining density estimation and a log scale automatically. It is common and reasonable, when plotting a graph with plasmid length on one axis, to use a log scale, because of the large range of plasmid lengths. However, “plotting a density on a log scale” could mean one of two very different things depending on the order that the density estimation and the log transformation are done. If the density estimation is done first, on the raw lengths, and then this estimate is plotted with a log-scaled axis, the result is like Figure 1B, which is a graph of the distribution of lengths (the numerical values of the plotted functions are the densities of plasmid lengths), although an unconventional one. But if the log transformation is done first, and then the density estimated from the log-transformed data, one gets an estimate of the distribution of log-lengths. Since Figure 1B is almost never what is actually wanted (it is hard to interpret and puts excessive emphasis on the lower tail of the distribution), graphing software asked to estimate a density and use a log scale will almost always choose to do the log transformation first.

Irrespective of the distribution of plasmid lengths, plasmid length and the biology of plasmids are closely entangled. For example, it is fairly clear from Figure 1 that there is a lower bound on the size of a conjugative plasmid (presumably set by the size of the genes needed to code for the complete conjugation machinery), and below this bound there are only mobilizable and nontransmissible plasmids (see also Coluzzi et al. 2022), while above it there are plasmids of all mobility classes. Another example is the decrease in the rate of transformation with plasmid size (Hanahan 1983). Plasmid size also influences the routes to adaptation: small plasmids are more likely to escape from bacterial restriction-modification systems through mutations in the target sequence than large plasmids, which due to their length contain more copies of the target sequence and need to resort to other means to escape restriction-modification systems (Shaw et al. 2023). Conversely, plasmid size may itself be the result of adaptation. Humphrey et al. (2021) hypothesize that the two modes in the size distribution of plasmids in *Staphylococcus aureus* (observed there on a linear scale; see also our Figure 3F) have evolved because those sizes are optimal for transduction by phages and *Staphylococcus aureus* pathogenicity islands (SaPis). It has further been found that small and large plasmids in *Klebsiella, Escherichia*, and *Salmonella*, as distinguished by the antimode of the log-length distribution, belong to different plasmid taxonomic units (Tal Dagan, private communication). It is possible that there is a fundamental reason why plasmids would cluster into plasmid taxonomic units by log-length, but it it also possible that the antimodes of the length and the log-length distributions are similar enough that the choice of distribution does not make a meaningful difference, i.e., both lead to nearly identical sets of small and large plasmids. Generally, since plasmid sizes and their distribution vary across bacterial families, within-family distributions of plasmid length are for many questions more informative than the distribution pooled across all plasmids.

The division of plasmids into large and small is said to be associated with a corresponding division into low and high copy number plasmids. It would be interesting to see if the distribution of plasmid copy numbers is bimodal, and to measure the extent to which it is associated with length. Unfortunately, it would be difficult to perform a similar analysis of the copy number distribution: plasmid copy number is not as frequently measured and reported as length, and is not included in databases of plasmid sequences. Moreover, the copy number is inherently more plastic than length, and may vary at different points of the host cell cycle or in different growth conditions, so that it may become impossible to speak of “the” copy number or to compare copy numbers of plasmids measured in different conditions.

To conclude, a clear understanding of the biology of plasmid length would be best served by clearly distinguishing between the distribution of plasmid lengths and that of log-lengths when discussing plasmid length, and presenting whichever is relevant to the question being asked. Since the fact that the two distributions have different shapes is not immediately apparent, confounding the two distributions can cause confusion or be misleading.

## Supporting information

R code for figures

R code for supplementary figures

## Acknowledgments

We thank Álvaro San Millán, Dustin Hanke, and Tal Dagan for helpful discussions and four anonymous reviewers for their comments that helped to improve the manuscript.

## Code availability

The R programs used to generate the figures in the text and the supplement are available as supplementary material.

