## Supplementary figures for "Is the distribution of plasmid lengths bimodal?"

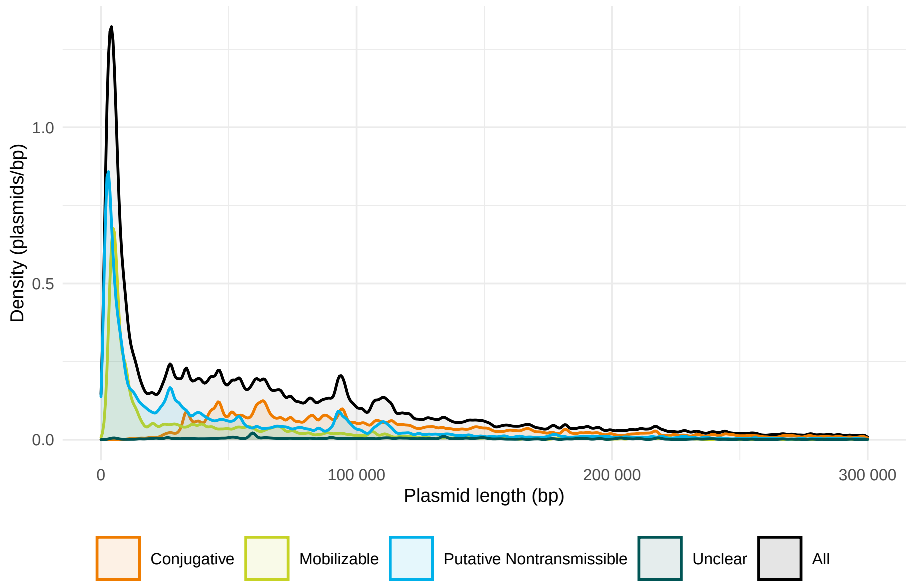


Figure S1: The distribution of plasmid lengths for plasmids shorter than 300 000 bp: in black, all plasmids; in colour, plasmids separated by mobility. The data sources and figure preparation are described in the caption to Figure 1 of the main text. The kernel density estimates are made with a bandwidth of 1000 bp, considerably smaller than the value of 6822.532 bp used for Figure 1A, to exclude the possibility of oversmoothing (the value used in Figure 1A is the median of the values chosen for each of the conjugative, mobilizable, and nontransmissible classes by R's default density estimation: this was done to ensure that all the densities in the figure were estimated with a common bandwidth). This estimate is, as expected, notably less smooth than the estimate in the main text, but it retains the same general features; in particular, it resembles the log-length distribution even less if anything, and it does not show particular signs of being bimodal.

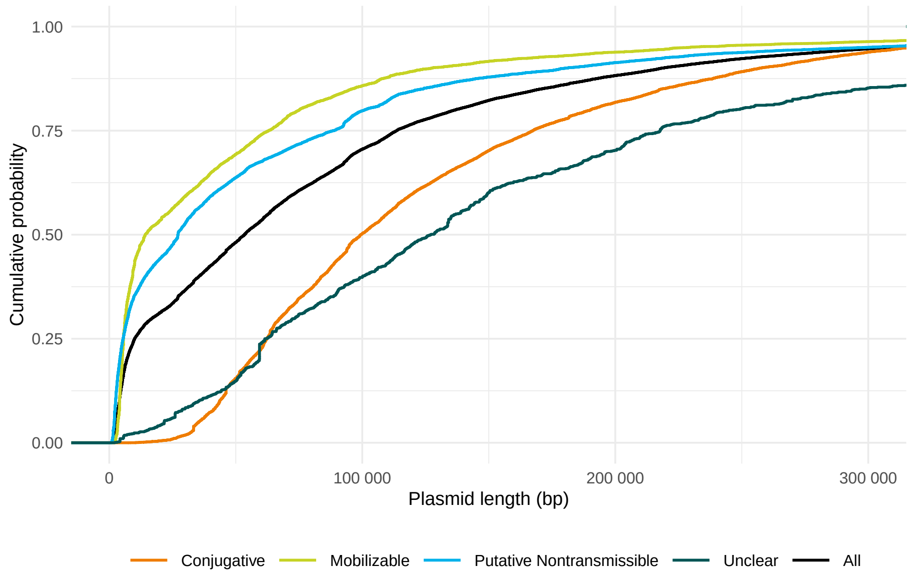


Figure S2: Empirical cumulative distribution function of the distribution of plasmid lengths: in black, all plasmids; in colour, plasmids separated by mobility. The data sources are described in the caption to Figure 1 in the main text. The cumulative distribution function has the advantage that there is no parameter choice involved in its estimation (unlike the need to choose a bandwidth for kernel density estimation or a bin width for a histogram), but is more difficult to interpret. A bimodal distribution should show two regions of steep slope separated by a region of shallow slope: there is no notably bimodal signs in the overall distribution. Note that unlike the kernel density estimates, each cumulative distribution function is normalized independently to a maximum of one.

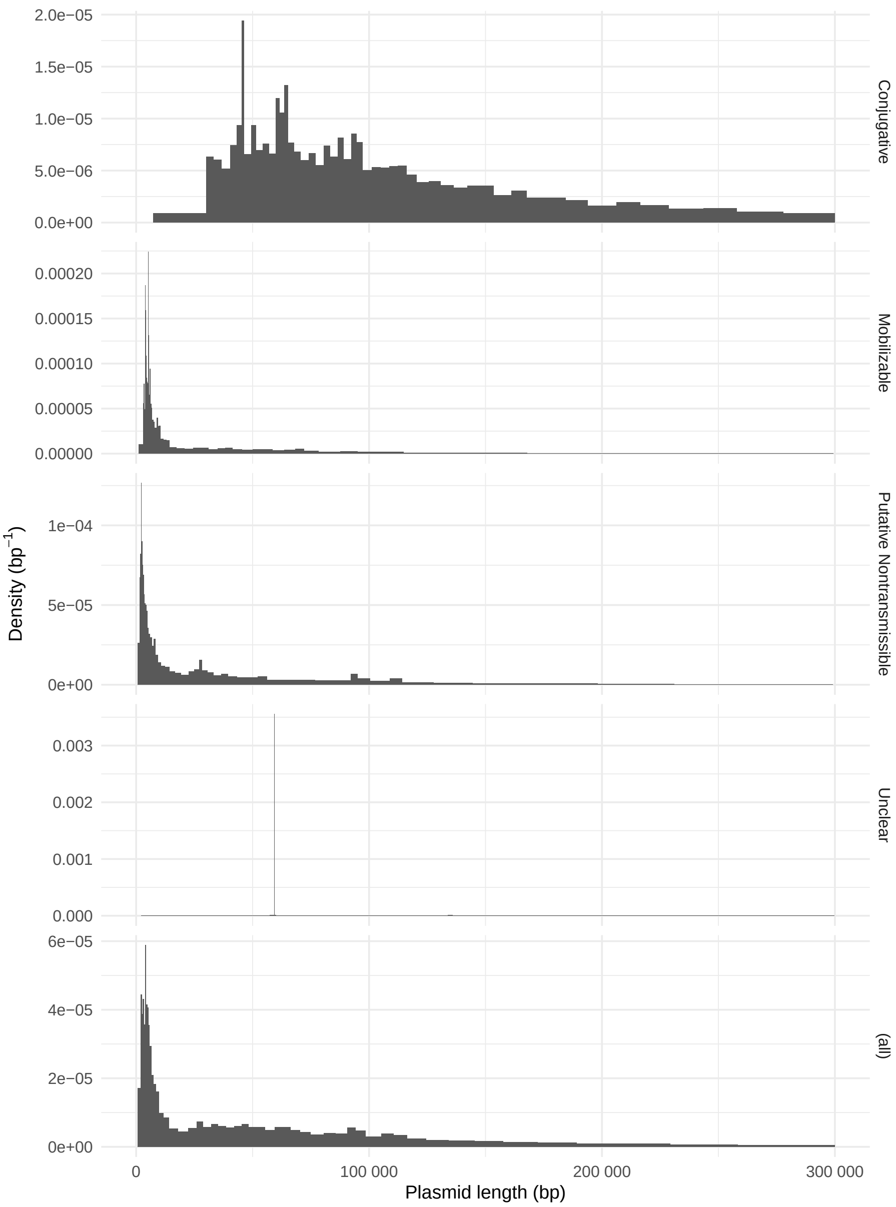


Figure S3: Equal-area histograms (percentograms) of the lengths of plasmids shorter than 300 000 bp. The data sources and figure preparation are described in the caption to Figure 1 of the main text. Each bin of the histogram holds 2 % of the total number of plasmids (i.e. the borders between bins are at the second, fourth, sixth,. . . percentiles). Yet again, the distribution looks notably different from the log-length distribution and the overall distribution is not particularly bimodal. Note that unlike the kernel density estimates, each percentogram is individually normalized to a total area of one.

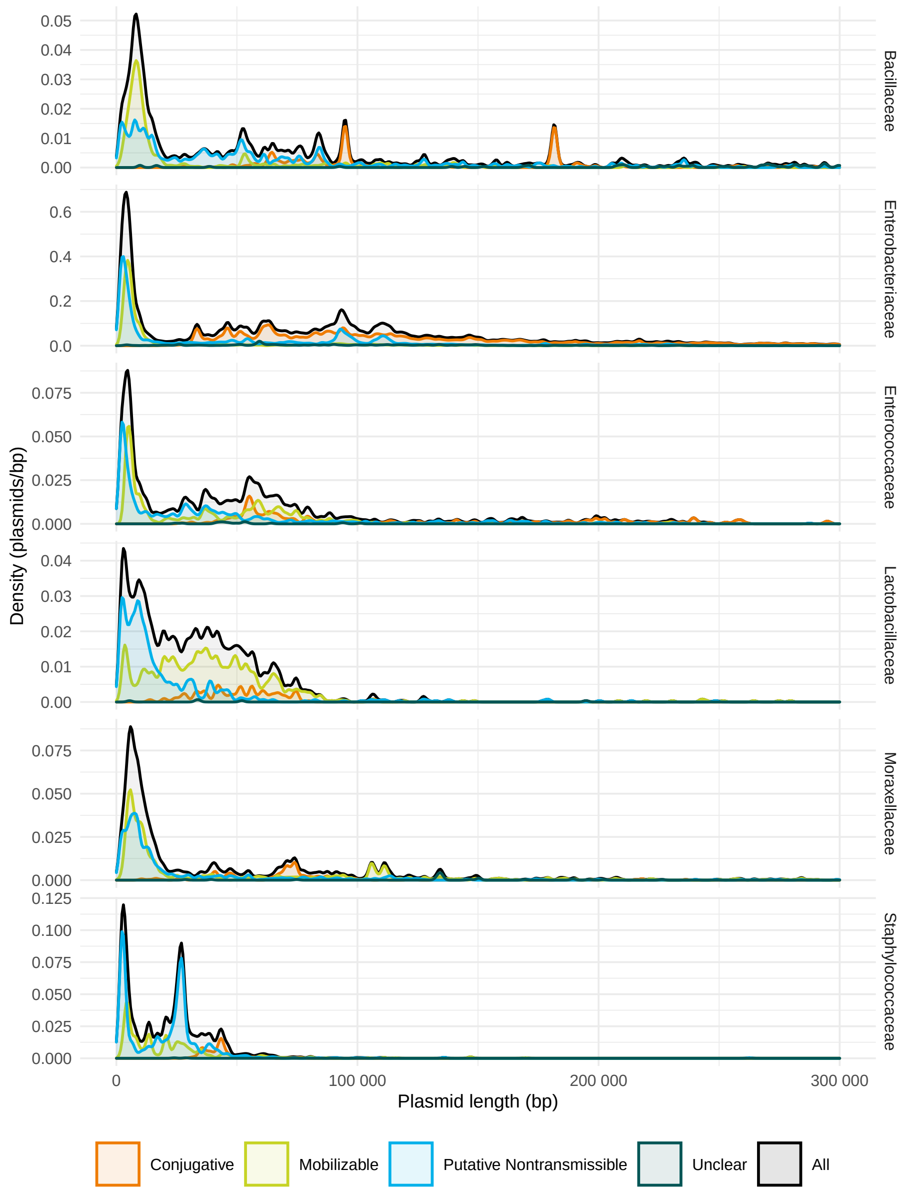


Figure S4: The distributions of plasmid lengths for plasmids shorter than 300 000 bp from the six bacterial families with at least 1000 plasmid sequences in the database: in black, all plasmids; in colour, plasmids separated by mobility. The data sources and figure preparation are described in the caption to Figure 1 of the main text. The kernel density estimates are made with a bandwidth of 1000 bp, considerably smaller than the values used in Figure 3 of the main text (which range from 4018.741 bp to 13 284.687 bp), to exclude the possibility of oversmoothing (the value used in each panel of Figure 3 is the median of the values chosen for each of the conjugative, mobilizable, and nontransmissible classes for that panel by R's default density estimation: this was done to ensure that all the densities in each panel were estimated with a common bandwidth).

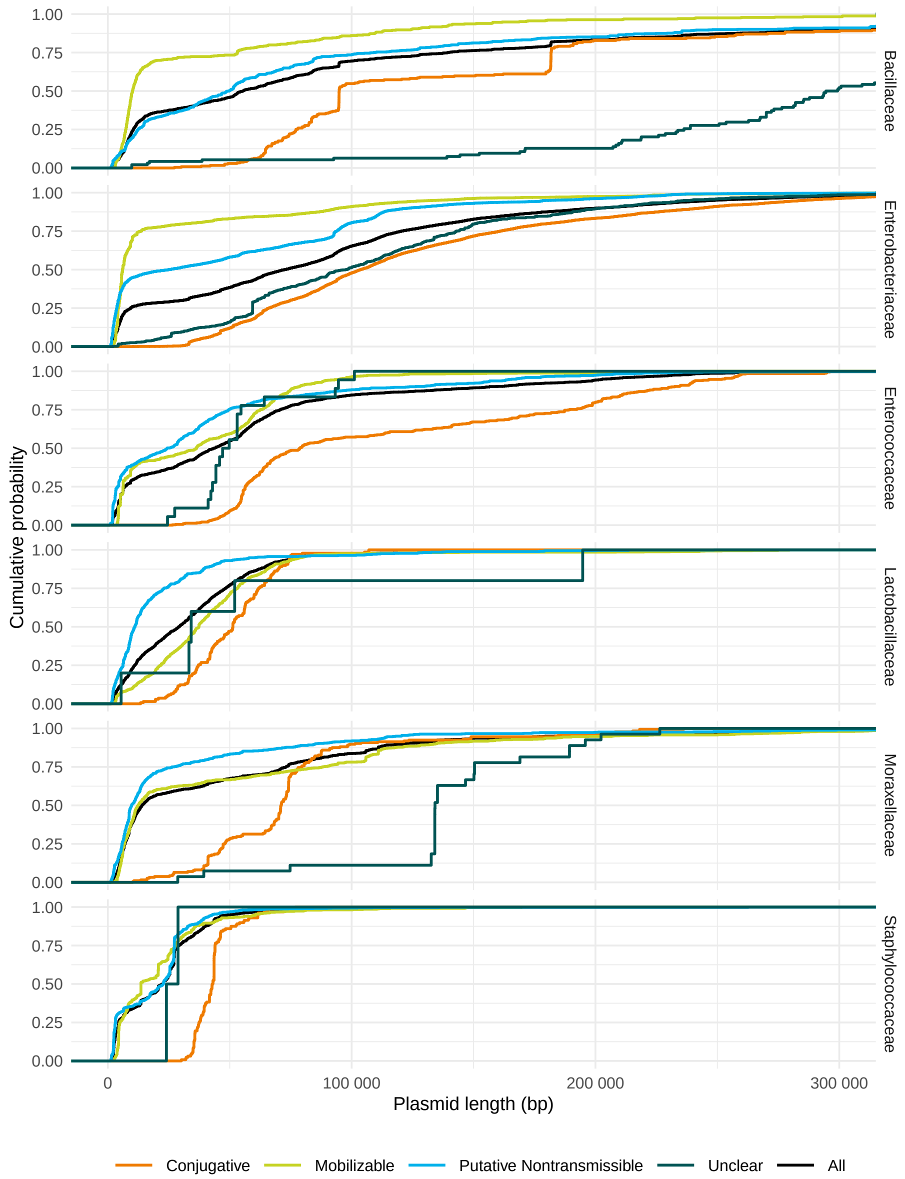


Figure S5: Empirical cumulative distribution functions of the distributions of plasmid lengths for plasmids from the six bacterial families with at least 1000 plasmid sequences in the database: in black, all plasmids; in colour, plasmids separated by mobility. The data sources are described in the caption to Figure 1 in the main text. A bimodal distribution should show two regions of steep slope separated by a region of shallow slope: many but not all families show a clear bimodal distribution. Note that unlike the kernel density estimates, each cumulative distribution function is normalized individually to a maximum of one.
