## Supplementary material for "Is the distribution of plasmid lengths bimodal?": R code for figures

library(tibble)
library(readr)
library(ggplot2)
library(gridExtra)
library(scales)

# The plasmid data is read from the file 'plsdb.tsv' in the archive
# available from
# https://ccb-microbe.cs.uni-saarland.de/plsdb/plasmids/download/plasmids_meta.tar.bz2
dat <- read_tsv('plsdb.tsv', col_types=cols_only(
	UID_NUCCORE=col_character(),
	Length_NUCCORE=col_integer(),
	taxon_family_name=col_character(),
	`relaxase_type(s)`=col_character(),
	mpf_type=col_character()))

# This drops three sequences (NZ_CP053353.1, NZ_CP030101.1, and
# NZ_CP069303.1) which are in PLSDB, but which are in fact chromosomal
# sequences. Of course there may be many other misidentified sequences,
# but we've chosen to remove just these since they are noticable by
# virtue of being the longest sequences.
dat <- dat[dat$Length_NUCCORE < 2800000,]

dat$Type <- factor(ifelse(is.na(dat$`relaxase_type(s)`),
	ifelse(is.na(dat$mpf_type), "Putative Nontransmissible",
		"Unclear"),
	ifelse(is.na(dat$mpf_type), "Mobilizable", "Conjugative")))
levels(dat$Type) <- c(levels(dat$Type), "All")
# "All" is a dummy level, which exists only to appear in the legend.

# We don't actually use this anywhere, but it prevents annoying errors
# from NAs.
dat$taxon_family_name[is.na(dat$taxon_family_name)] <- "Uncultured"

# We have to do the bandwidth selection for the kernel density estimates
# somewhat manually. We are plotting a density for each group, and then
# the density for all plasmids, which should be the sum of the group
# densities. But ggplot calculates the bandwidth separately for each
# group, so they are not compatible; in some cases there can even be
# artifacts where a group density exceeds the total density. Therefore,
# we have to choose a single bandwidth for each panel of a plot, to be
# used for all of the group densities and the total density; the
# selected bandwidth is the median of the bandwidths for the
# conjugative, mobilizable, and nontransmissible subgroups, which in
# practice seems to mean that of the nontransmissible subgroup (we
# exclude unclear plasmids because they're weird). We still use bw.nrd0
# here instead of switching to the allegedly superior bw.SJ because I
# still want to stay close to the default settings of R.
bw.lin <- median(c(
	bw.nrd0(dat$Length_NUCCORE[dat$Type == "Conjugative"]),
	bw.nrd0(dat$Length_NUCCORE[dat$Type == "Mobilizable"]),
	bw.nrd0(dat$Length_NUCCORE[
		dat$Type == "Putative Nontransmissible"])))
bw.log <- median(c(
	bw.nrd0(log10(dat$Length_NUCCORE[dat$Type == "Conjugative"])),
	bw.nrd0(log10(dat$Length_NUCCORE[dat$Type == "Mobilizable"])),
	bw.nrd0(log10(dat$Length_NUCCORE[
		dat$Type == "Putative Nontransmissible"]))))
# Determining the KDE bandwidth gets particularly annoying with the "by
# family" plot, since we want to choose a separate bandwidth for each
# facet and ggplot doesn't treat bw as an aesthetic. We'll use the
# following function to do so.
bw.family <- function(family, logged) {
	res <- c()
	ts <- c("Conjugative","Mobilizable","Putative Nontransmissible")
	for (t in ts) {
		if (logged) {
			x <- log10(dat$Length_NUCCORE[
				dat$taxon_family_name == family &
				dat$Type == t])
		} else {
			x <- dat$Length_NUCCORE[
				dat$taxon_family_name == family &
				dat$Type == t]
		}

		if (length(x) > 1) {
			res <- c(res, bw.nrd0(x))
		}
	}
	median(res)
}

f <- function(x) g(log(x)) / x
g <- function(x) (dnorm(x,mean=0.1) + dnorm(x,mean=2.9)) / 2

focal.families <- c("Bacillaceae","Enterobacteriaceae",
	"Enterococcaceae","Lactobacillaceae","Moraxellaceae",
	"Staphylococcaceae")
fams <- tibble(taxon_family_name=rep(focal.families, 2),
	logged=c(rep(FALSE, length(focal.families)),
		rep(TRUE, length(focal.families))),
	antimode=c(NA,3.0e4,2.8e4,2.7e4,NA,1.4e4,2.4e4,1.9e4,
		1.5e4,1.6e4,2.7e4,8.5e3),
	nlab="",
	tag=LETTERS[1:(2*length(focal.families))])
for (fam in focal.families)
	fams[fams$taxon_family_name == fam,"nlab"] <-
		paste("n =", nrow(dat[dat$taxon_family_name == fam,]))

# Extract the legend from an "idealized version" of the plots we will be
# showing. This is needed for two reasons:
# - to allow us to arrange the legend in any position relative to the
#   panels of the figure
# - to prevent ggplot from getting the colours wrong, since it gets
#   confused by us plotting the total density and densities per plasmid
#   type together.
# The disadvantage is that we now need to keep the legend colours and
# the colours on the plots in sync manually.
# The code to do this is based on a StackOverflow answer I found.
options(ggplot2.discrete.colour=c("#ef7c00", "#c6d325", "#00b1ea",
	"#005555", "#000000"))
tmp <- ggplot_gtable(ggplot_build(
	ggplot(dat, aes(x=Length_NUCCORE)) +
	geom_density(aes(fill=Type, colour=Type),alpha=0.1) +
	scale_colour_discrete(drop=FALSE, guide=guide_legend(nrow=2)) +
	scale_fill_discrete(drop=FALSE, guide=guide_legend(nrow=2)) +
	theme_minimal(base_size=7) +
	theme(legend.position="bottom",
		legend.title=element_blank())))$grobs
legend <- tmp[[which(sapply(tmp, function(x) x$name) == "guide-box")]]

plt1a <- ggplot(dat, aes(x=Length_NUCCORE,y=after_stat(count))) +
	geom_density(bw=bw.lin,n=2048,alpha=0.05,fill="black") +
	geom_density(aes(fill=Type,color=Type),
		bw=bw.lin,n=2048,alpha=0.1) +
	scale_x_continuous(labels=label_number()) +
	coord_cartesian(xlim=c(0,3e5)) +
	labs(x="Plasmid length (bp)",
		y="Density (plasmids/bp)",
		tag="A") +
	theme_minimal(base_size=7) +
	theme(legend.position="none", plot.tag.position="topright")
plt1b <- ggplot(dat, aes(x=Length_NUCCORE,y=after_stat(count))) +
	geom_density(bw=bw.lin,n=2**16) +
	geom_density(aes(color=Type), bw=bw.lin, n=2**16) +
	scale_x_continuous(breaks=c(1e3,1e4,1e5,1e6),
		labels=expression(10^3,10^4,10^5,10^6)) +
	coord_trans(x="log10") +
	labs(x="Plasmid length (bp, log scale)",
		y="Density (plasmids/bp)",
		tag="B") +
	theme_minimal(base_size=7) +
	theme(legend.position="none", plot.tag.position="topright")
plt1c <- ggplot(dat, aes(x=Length_NUCCORE,y=after_stat(count))) +
	geom_density(bw=bw.log,alpha=0.05,fill="black") +
	geom_density(aes(fill=Type,color=Type), bw=bw.log, alpha=0.1) +
	scale_x_log10(labels=trans_format("log10", math_format(.x))) +
	labs(x="Log plasmid length (ban)",
		y="Density (plasmids/ban)",
		tag="C") +
	theme_minimal(base_size=7) +
	theme(legend.position="none", plot.tag.position="topright")
ggsave('letter-fig-distribution.pdf',
	arrangeGrob(plt1a,plt1b,plt1c,legend,nrow=4,heights=c(6,6,6,2)),
	width=90, height=150, unit="mm")
plt2a <- ggplot(data.frame(x=c(0,25)),aes(x)) +
	stat_function(fun=f,n=501,geom="line") +
	stat_function(fun=f,n=501,geom="area",alpha=0.1) +
	labs(x="s", y="Probability density f(s)", tag="X") +
	theme_minimal(base_size=7) +
	theme(plot.tag.position="topright")
plt2b <- ggplot(data.frame(x=c(-4,6)),aes(x)) +
	stat_function(fun=g,geom="line") +
	stat_function(fun=g,geom="area",alpha=0.1) +
	labs(x="t", y="Probability density g(t)", tag="log X") +
	theme_minimal(base_size=7) +
	theme(plot.tag.position="topright")
ggsave('letter-fig-analytic.pdf',
	arrangeGrob(plt2a,plt2b,nrow=1,widths=c(9,11)),
	width=90, height=60, unit="mm")
plt3a <- ggplot(NULL, aes(x=Length_NUCCORE,y=after_stat(count))) +
	lapply(focal.families, function(fam)
		geom_density(data=dat[dat$taxon_family_name == fam,],
			bw=bw.family(fam,FALSE),
			n=2048,
			alpha=0.05,
			fill="black")) +
	lapply(focal.families, function(fam)
		geom_density(aes(fill=Type,color=Type),
			data=dat[dat$taxon_family_name == fam,],
			bw=bw.family(fam,FALSE),
			n=2048,
			alpha=0.1)) +
	geom_vline(data=fams[!fams$logged,],
		aes(xintercept=antimode),
		color="grey70",
		linetype=2) +
	geom_text(data=fams[!fams$logged,],
		aes(x=antimode,y=Inf,label=antimode),
		hjust=-0.1,
		vjust=1.1,
		color="grey70",
		size=2) +
	geom_text(data=fams[!fams$logged,],
		aes(x=Inf,y=Inf,label=tag),
		hjust=1.1,
		vjust=1.1,
		size=3) +
	facet_grid(rows=vars(taxon_family_name), scales="free_y") +
	scale_x_continuous(labels=label_number()) +
	coord_cartesian(xlim=c(0,3e5)) +
	scale_y_continuous(expand=expansion(mult=c(0,0.05))) +
	labs(x="Plasmid length (bp)", y="Density (plasmids/bp)") +
	theme_minimal(base_size=7) +
	theme(legend.position="none", strip.text=element_blank())
plt3b <- ggplot(NULL, aes(x=Length_NUCCORE,y=after_stat(count))) +
	lapply(focal.families, function(fam)
		geom_density(data=dat[dat$taxon_family_name == fam,],
			bw=bw.family(fam,TRUE),
			alpha=0.05,
			fill="black")) +
	lapply(focal.families, function(fam)
		geom_density(aes(fill=Type,color=Type),
			data=dat[dat$taxon_family_name == fam,],
			bw=bw.family(fam,TRUE),
			alpha=0.1)) +
	geom_vline(data=fams[fams$logged,],
		aes(xintercept=antimode),
		color="grey70",
		linetype=2) +
	geom_text(data=fams[fams$logged,],
		aes(x=antimode,y=Inf,label=antimode),
		hjust=1.1,
		vjust=1.1,
		color="grey70",
		size=2) +
	geom_text(data=fams[fams$logged,],
		aes(x=Inf, y=Inf, label=nlab),
		hjust=1.1,
		vjust=3.0,
		color="grey70",
		size=2) +
	geom_text(data=fams[fams$logged,],
		aes(x=Inf,y=Inf,label=tag),
		hjust=1.1,
		vjust=1.1,
		size=3) +
	facet_grid(rows=vars(taxon_family_name), scales="free_y") +
	scale_x_log10(labels=trans_format("log10", math_format(.x))) +
	scale_y_continuous(expand=expansion(mult=c(0,0.05))) +
	labs(x="Log plasmid length (ban)", y="Density (plasmids/ban)") +
	theme_minimal(base_size=7) +
	theme(legend.position="none")
ggsave('letter-fig-family.pdf',
	arrangeGrob(arrangeGrob(plt3a,plt3b,nrow=1,widths=c(11,12)),
		legend,
		nrow=2,heights=c(10,1)),
	width=140, height=180, units="mm")
