## Supplementary material for "Is the distribution of plasmid lengths bimodal?": R code for supplementary figures

library(tibble)
library(readr)
library(ggpercentogram)
library(ggplot2)
library(gridExtra)
library(scales)

# The plasmid data is read from the file 'plsdb.tsv' in the archive
# available from
# https://ccb-microbe.cs.uni-saarland.de/plsdb/plasmids/download/plasmids_meta.tar.bz2
dat <- read_tsv('plsdb.tsv', col_types=cols_only(
	UID_NUCCORE=col_character(),
	Length_NUCCORE=col_integer(),
	taxon_family_name=col_character(),
	`relaxase_type(s)`=col_character(),
	mpf_type=col_character()))
# This drops three sequences (NZ_CP053353.1, NZ_CP030101.1, and
# NZ_CP069303.1) which are in PLSDB, but which are in fact chromosomal
# sequences. Of course there may be many other misidentified sequences,
# but we've chosen to remove just these since they are noticable by
# virtue of being the longest sequences.
dat <- dat[dat$Length_NUCCORE < 2800000,]

dat$Type <- factor(ifelse(is.na(dat$`relaxase_type(s)`),
	ifelse(is.na(dat$mpf_type), "Putative Nontransmissible", "Unclear"),
	ifelse(is.na(dat$mpf_type), "Mobilizable", "Conjugative")))
levels(dat$Type) <- c(levels(dat$Type), "All")
# "All" is a dummy level, which exists only to appear in the legend.

focal.families <- c("Bacillaceae", "Lactobacillaceae", "Moraxellaceae",
	"Staphylococcaceae", "Enterococcaceae", "Enterobacteriaceae")
dat.fam <- dat[dat$taxon_family_name %in% focal.families,]

# Extract the legend from an "idealized version" of the plots we will be
# showing. This is needed for two reasons:
# - to allow us to arrange the legend in any position relative to the
#   panels of the figure
# - to prevent ggplot from getting the colours wrong, since it gets
#   confused by us plotting the total density and densities per plasmid
#   type together.
# The disadvantage is that we now need to keep the legend colours and
# the colours on the plots in sync manually.
# The code to do this is based on a StackOverflow answer I found.
options(ggplot2.discrete.colour=c("#ef7c00", "#c6d325", "#00b1ea",
	"#005555", "#000000"))
tmp <- ggplot_gtable(ggplot_build(
	ggplot(dat, aes(x=Length_NUCCORE)) +
	geom_density(aes(fill=Type, colour=Type),alpha=0.1) +
	scale_colour_discrete(drop=FALSE) +
	scale_fill_discrete(drop=FALSE) +
	theme_minimal(base_size=7) +
	theme(legend.position="bottom",
		legend.title=element_blank())))$grobs
legend <- tmp[[which(sapply(tmp, function(x) x$name) == "guide-box")]]

plt1 <- ggplot(dat, aes(x=Length_NUCCORE,y=after_stat(count))) +
	geom_density(bw=1000,n=1024,alpha=0.05,fill="black") +
	geom_density(aes(fill=Type,color=Type), bw=1000, alpha=0.1) +
	scale_x_continuous(labels=label_number()) +
	coord_cartesian(xlim=c(0,3e5)) +
	labs(x="Plasmid length (bp)", y="Density (plasmids/bp)") +
	theme_minimal(base_size=7) +
	theme(legend.position="none")
ggsave('letter-supp-fig-constbw.pdf',
	arrangeGrob(plt1,legend,heights=c(6,1)),
	width=121, height=80, units="mm")
plt2 <- ggplot(dat, aes(x=Length_NUCCORE)) +
	stat_ecdf(geom="step") +
	stat_ecdf(aes(color=Type), geom="step") +
	scale_x_continuous(labels=label_number()) +
	scale_colour_discrete(drop=FALSE) +
	coord_cartesian(xlim=c(0,3e5)) +
	labs(x="Plasmid length (bp)", y="Cumulative probability") +
	theme_minimal(base_size=7) +
	theme(legend.position="bottom", legend.title=element_blank())
ggsave('letter-supp-fig-ecdf.pdf', plt2,
	width=121, height=80, units="mm")
plt3 <- ggplot(dat, aes(x=Length_NUCCORE)) +
	geom_percentogram(bins=50) +
	scale_x_continuous(labels=label_number()) +
	coord_cartesian(xlim=c(0,3e5)) +
	facet_grid(rows=vars(Type), margins=TRUE, scales="free_y") +
	labs(x="Plasmid length (bp)",
		y=expression(paste("Density (", plain(bp)^{-1}, ")"))) +
	theme_minimal(base_size=7) +
	theme(legend.position="none")
ggsave('letter-supp-fig-percentogram.pdf', plt3,
	width=121, height=160, units="mm")
plt5 <- ggplot(dat.fam, aes(x=Length_NUCCORE,y=after_stat(count))) +
	geom_density(bw=1000,n=1024,alpha=0.05,fill="black") +
	geom_density(aes(fill=Type,color=Type), bw=1000, alpha=0.1) +
	scale_x_continuous(labels=label_number()) +
	coord_cartesian(xlim=c(0,3e5)) +
	facet_grid(rows=vars(taxon_family_name), scales="free_y") +
	labs(x="Plasmid length (bp)", y="Density (plasmids/bp)") +
	theme_minimal(base_size=7) +
	theme(legend.position="none")
ggsave('letter-supp-fig-fam-constbw.pdf',
	arrangeGrob(plt5,legend,heights=c(13,1)),
	width=121, height=160, units="mm")
plt6 <- ggplot(dat.fam, aes(x=Length_NUCCORE)) +
	stat_ecdf(geom="step") +
	stat_ecdf(aes(color=Type), geom="step") +
	scale_x_continuous(labels=label_number()) +
	scale_colour_discrete(drop=FALSE) +
	coord_cartesian(xlim=c(0,3e5)) +
	facet_grid(rows=vars(taxon_family_name), scales="free_y") +
	labs(x="Plasmid length (bp)", y="Cumulative probability") +
	theme_minimal(base_size=7) +
	theme(legend.position="bottom", legend.title=element_blank())
ggsave('letter-supp-fig-fam-ecdf.pdf', plt6,
	width=121, height=160, units="mm")
